# Decoding brain-wide signatures of uninformed choices for BCI assisted decision-making

**DOI:** 10.1101/2025.11.12.688013

**Authors:** Sophia Gimple, Maxime Verwoert, Laura Marras, Paul Weger, Steffen A. Herff, Johannes P. van Dijk, Simon Tousseyn, Pieter L. Kubben, Yasin Temel, Marcus L.F. Janssen, Christian Herff

## Abstract

Decision-making is an essential cognitive function. It can be impaired due to a number of neurological and psychiatric disorders as well as external factors such as time pressure or stress. To assist users during decision-making, we propose a decision-making brain computer interface (BCI) that can alert to uninformed decision-making, prompt additional information seeking and therefore improve decision-making quality. To this aim, we establish the feasibility of decoding uninformed decision-making from local field potentials recorded with implanted stereo-electroencephalography (sEEG) electrodes in 6 participants. We show that decoding of available information above chance level is possible for all participants, both after stimulus presentation, as well as before task response. Starting from stimulus onset, the temporal processing hierarchy of informed vs. uninformed decision-making spans from visual processing through hippocampal memory processes to frontal control network shifting. The anterior insula, known to be a decision-making hub, codes available information during the decision phase prior to button press. These results further elucidate the neural basis of coded information availability and confirm the feasibility of a decision-making BCI.

## Introduction

Decision-making is a cognitive process fundamental both in every-day situations, such as choosing which clothes to wear, as well as in high-stake situations like making a big financial investment. In all situations, the success in decision-making can be characterized as a trade-off between speed and accuracy^1^. While time is a limited resource and delaying decisions can make the decision itself irrelevant, incorrect decisions can lead to suboptimal decision outcomes with potentially serious consequences^1^. This relationship can also be directly demonstrated in simple decision-making paradigms, in which inducing time pressure leads to increased error rates^2^. Sequential sampling models propose that when making a decision, information is collected from the environment until an internal threshold of certainty is reached^3^. Previous studies, however, have shown that humans self-judgment of decision confidence does not accurately reflect the foundation of available information during decision-making^4^. In the extremes that can lead to a very confident uninformed decision of likely low quality or the delay of a decision due to low confidence despite a solid informational basis. This effect can be increased due to external factors such as stress, emotions or sleep deprivation that can further impact decision quality^5–7^. Furthermore, in a variety of psychiatric, developmental and neurodegenerative diseases such as depression, anxiety, bipolar disorders, Parkinson’s disease, and ADHD, decision-making is also often effected e.g. through impulsivity or indecisiveness^8–11^. At the same time, we live in an age when large amounts of information are readily accessible at any time from a variety of sources. Therefore, seeking of additional information can be easily implemented during the decision-making process to improve decision accuracy where necessary^12,13^. To bridge this gap between the possibility of quick information search and suboptimal judgment of how informed a decision is, not accurately represented by decision confidence, we envision a cognitive decision-making brain computer interface (BCI). A cognitive BCI^14^ can serve as a valuable tool for measuring cognitive states to directly adapt a human-computer interface or alert the user and their surroundings. More specifically, we propose a BCI able to objectively measure subconscious information availability and prompt users to optimize their decision-making behavior. Such an interface could help with improving decision quality both in a general public as well as for user populations making high-stake decisions, being exposed to factors impairing decision-making competence or patient-populations.

Neurobiologically based models split decision-making into different subprocesses. Ernst and Paulus^15^ define the key processes as (1) building preferences between different options, (2) the selection of one option and lastly (3) feedback evaluation. They link those separate processes of decision-making to several relevant brain regions including dorsolateral prefrontal cortex (DLPFC), dorsal anterior cingulate cortex (dACC), presupplementary motor area (preSMA), anterior Insula, Amygdala and dorsal Striatum. Previous research identifies a dedicated neural decision variable representing the temporal accumulation of signals until a decision between competing options is reached, both in monkeys^16^ and humans^17^. Additionally, prior studies show that decision-making outcome can be decoded from a number of brain regions, such as premotor and motor cortex^16^, parietal cortex^18^, temporal cortex^19^, orbitalfrontal cortex^20^, prefrontal cortex, Insula and anterior cingulate cortex (ACC)^21^. This implies the significance of neural processes in these brain regions for decision option selection in different decision-making situations and tasks. On top of that, other decision parameters can also be decoded from neural signals, as demonstrated in several studies discussed below. A study by Thiery et al. identifies neural processes that reflect whether a movement is instructed or a result of free choice^22^. In an EEG study with healthy participants deciding on which ear they are presented with more clicks, Kubanek et al. predict self-reported decision confidence from cortical alpha activity^23^. Similarly, Desender et al. identify stimulus locked post-decision event related potential (ERP) markers significantly modulated by confidence in a color classification task with seventeen participants. Furthermore, the authors show a link between the identified ERP markers and additional information seeking behavior^12^. In addition, Valeriani et al. use EEG features aligned to stimulus presentation to differentiate between conditions in multiple decision parameters^24^. These parameters include self-reported decision confidence which is proposed to be useful in a BCI for improvement of AI and human cooperation. They identify temporal neural markers utilizing low-frequency time-domain features in parietal, occipital and frontal-temporal electrodes and frontal-temporal alpha as a relevant oscillatory feature^24^. For the development of a decision-making BCI that can alert users to uninformed decision-making, we are however particularly interested in the neural coding of available information and more specifically the differentiation between uninformed and informed decision-making processes. Sadras and colleagues demonstrate the feasibility of decoding information availability from a down-sampled, low-frequency, stimulus-locked time-series that can potentially be implemented in an EEG-based decision-making BCI^25^. In their experimental design, participants are asked to determine whether a person is depicted wearing a cap or a helmet based on a picture. The available information during decision-making is modulated by blurring the pictures to varying degrees to create more and less difficult trials. For each of the 11 participants, high and low-information levels can be predicted from a low frequency time-domain feature with receiver operating characteristic area under the curve ((roc-) AUC) values above 0.5 with at least one of the evaluated classifiers. On average, decoding AUC reaches values of up to 0.75 using a logistic regression classifier during a time interval between 0.2 and 0.7s post stimulus-onset. It should be noted that using this approach, the authors can classify high- and low-information levels during decision-making time-locked to stimulus onset but not locked to participant response^25^. Even though this EEG approach seems very promising, previous literature establishes the importance of both cortical and subcortical regions for decision-making (e.g.^15^). We expand on previous research by evaluating the decodability of uninformed decisions from sEEG electrodes, which provide insights into the intricate decision-making processes from both surface and deep brain regions (compare Fig.1). To this aim, we first demonstrate discrimination between informed and uninformed decision-making, evaluating different neurophysiological features. In a second step, we establish regions of interest and time windows both locked to stimulus onset, as well as to participant response. With these analyses, we investigate the temporal dynamics of neural processes underlying decision-making that are linked to information availability during the decision. Our results could inform future attempts of creating BCIs that support users during decision-making, for example by prompting users to engage in additional information seeking behavior in the case of uninformed decision-making.

## Results

### Decision-making quality improved in informed trials

In a decision-making task, participants were asked to assign visual stimuli, each presented three times, to one of two classes (left or right) and were given feedback on their decision. Participants reached a higher decision-making performance during informed decision-making (third stimulus presentation, mean accuracy 0.76, std=0.12) compared to uninformed decision-making (first stimulus presentation, mean accuracy 0.42, std=0.14). This difference was significant (t(5)=−8.24, p*<*0.001), indicating that the participants were indeed able to learn class membership (left or right button press) of stimuli during previous presentations of the same stimulus (Fig. 1 d). Participants responded on average after 1.35 seconds (std=0.37, median=1.36) during the third stimulus presentation, compared to 1.45 seconds (std=0.40, median=1.40) during the first stimulus presentation (Fig. 1 a linear mixed effects model controlling for participant differences revealed that the response time differed significantly between trials where the stimulus was presented for the first, second or third time (b=−0.05, SE=0.02, z=−2.85, p*<*0.01, 95% CI=[−0.087, −0.016]). Post-hoc Tukey tests revealed that response time was significantly faster during third compared to first stimulus presentation (p=0.03). However, as also visible from Fig. 1 c) the difference in response time is not consistent across participants. While for some participants the difference in response time is clearly observable, for two participants the mean response time differences between first and third stimulus presentation trial are close to zero with one participant even reacting slower on average during the third stimulus presentation. Generally, prompts instructed participants to respond no earlier than 1 second after stimulus presentation. However, the 6 participants responded within the first second on average in 15 (std=16.11) out of the 90 trials.

**Figure 1.**
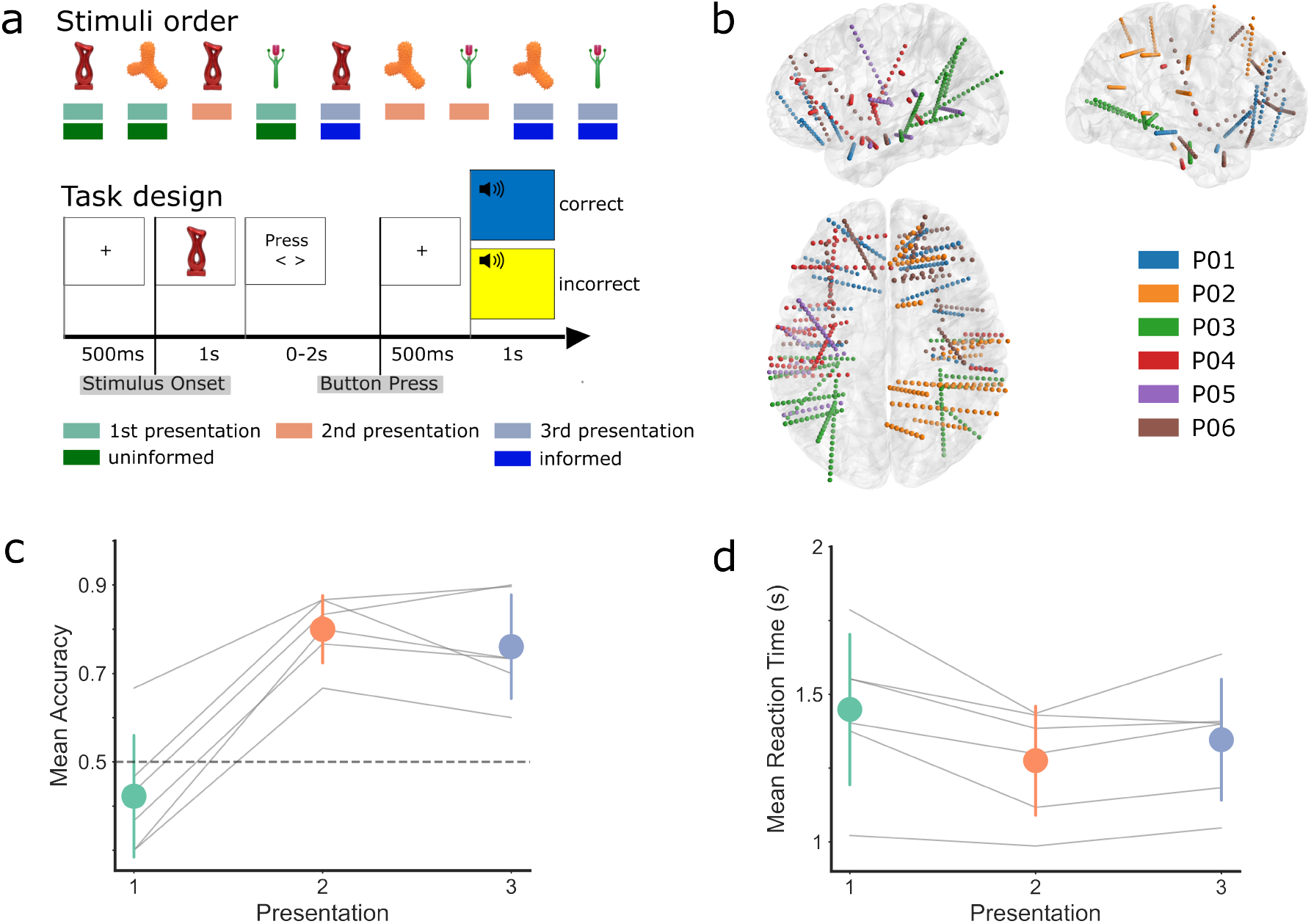
Task, setup and behavioral results. a) During one task block, three visual stimuli were presented three times in randomized order. After stimulus presentation, participants were asked to decide to which of two groups (left or right) the respective stimulus belonged, feedback was provided afterwards. For the first presentation of the stimulus, we consider the decision to be uninformed while for the third presentation, an informed decision based on previous feedback was assumed. b) Electrode locations of 6 participants visualized in an MNI brain show the wide coverage of both cortical and subcortical brain regions. c) Mean and standard deviation of task accuracy of all participants and mean accuracy per participant show the performance of the participants in assigning a stimulus to the correct group (left or right). The dashed line indicates the chance level at 50% expected during random guessing. d) Mean reaction times of all participants individually and mean and standard deviation of all participants are visualized for first, second and third stimulus presentation separately.

### Temporal dynamics of neural coding of uninformed decision-making after stimulus onset

Decoding informed versus uninformed decision-making aligned to stimulus onset is possible from multiple neural processes in different frequency bands and brain regions, as summarized in Fig. 2 and Fig. 3. Low-frequency filtered time-domain (LF-time) features with a mean prediction AUC of 79% (std=0.08; b=0.29, SE=0.03, z=8.47, p*<*0.0001, 95% CI=[0.23, 0.36]) and gamma frequency band features with a mean prediction AUC of 75% (std=0.06, b=0.25, SE=0.03, z=9.41, p*<*0.0001, 95% CI=[0.20, 0.30]) provided the best results when combining all channels (compare Fig. 2). Additionally, delta with a mean AUC of 65% (std=0.10; b=0.15, SE=0.05, z=3.17, p*<*0.001, 95% CI=[0.06, 0.24]) and beta features with a mean AUC of 62 % (std=0.07; b=0.12, SE=0.03, z=3.89, p*<*0.0001, 95% CI=[0.06, 0.18]) achieved significant decoding above chance level. The AUCs of theta with a mean of 55% (std=0.12; b=0.05, SE=0.06, z=0.90, p=n.s., 95% CI=[−0.06, 0.16]) and alpha frequency band features with mean 58% (std=0.13; b=0.08, SE=0.05, z=1.48,p=n.s., 95% CI=[−0.03, 0.19]) were not significantly above chance level (50%). When zooming in, on decodability of information levels from single regions of interest, we observed a clear temporal pattern visualized in Fig. 3. The earliest significant decodability was observed in the occipital-temporal sulcus when decoding from low frequency time domain features at around 400 ms after stimulus onset. Around 500 ms after stimulus onset, time-domain features in hippocampus and around 550ms white-matter gamma power features started showing significant decoding between informed and uninformed decision processes. Last, around 700 ms, delta power in frontal brain regions predicted a difference between informed and uninformed neural processes. Temporal features in occipital temporal sulcus and white matter gamma power showed a fast decrease in predictability between informed and uninformed decision-making within the first second after stimulus onset. Especially, usage of temporal features in hippocampus resulted in a prolonged consistent significant decodability with high AUC, exceeding 1 second and therefore potentially lasting until button press.

**Figure 2.**
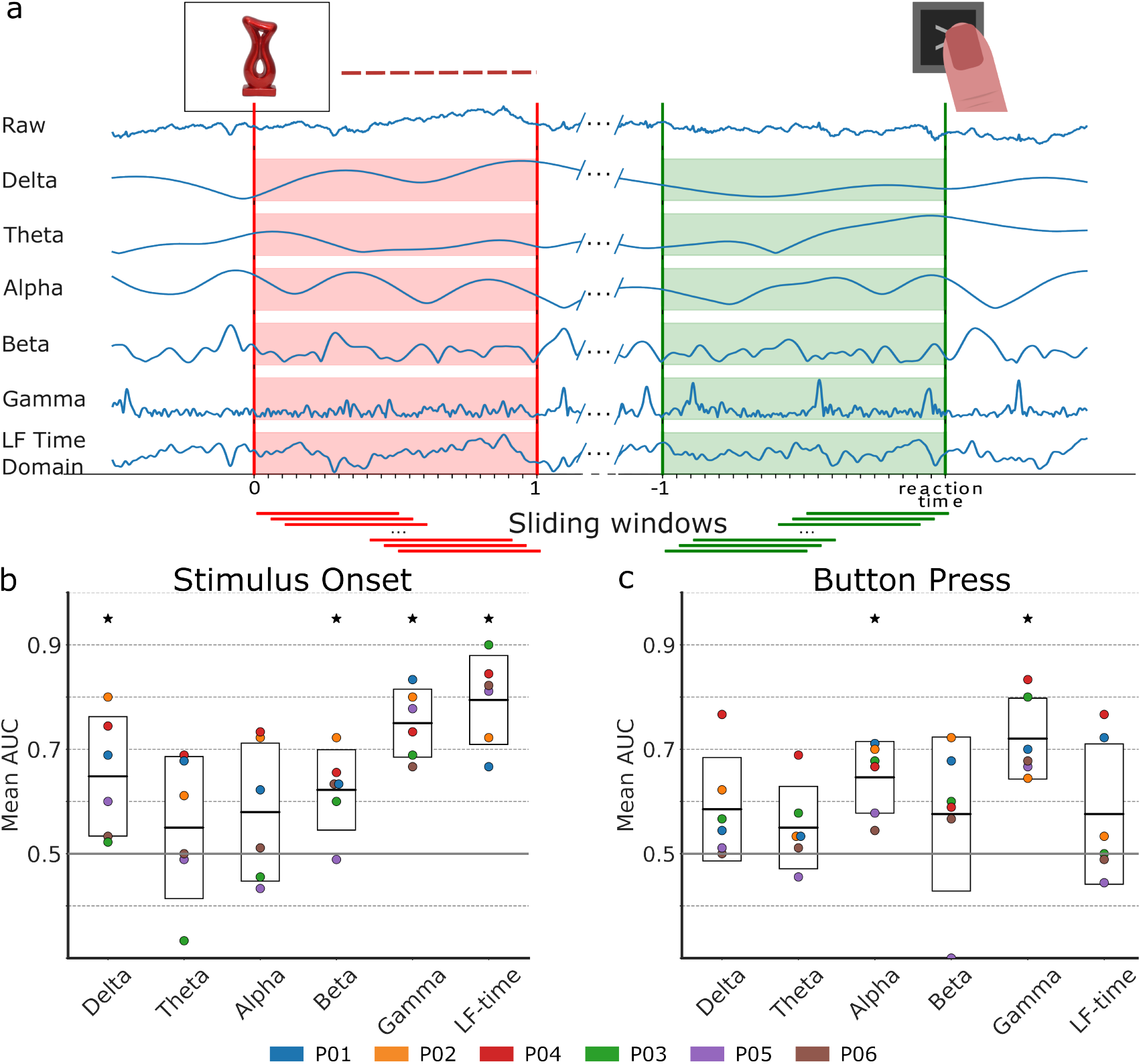
Analysis pipeline and decoding results a) From the raw neurophysiological signal five frequency band features and a time-domain signal were extracted. The data was split into time windows capturing 1 second after stimulus onset and 1 second before button press. b) Decoding results classifying between informed and uninformed decisions from different features from all time-windows and channels combined. Error boxes show mean and standard deviation of AUC across all participants combined. Mean AUC of all participants individually is visualized using colored dots. Significance is determined using a linear mixed-effects model with participant number as a random factor and indicated with a black star.

**Figure 3.**
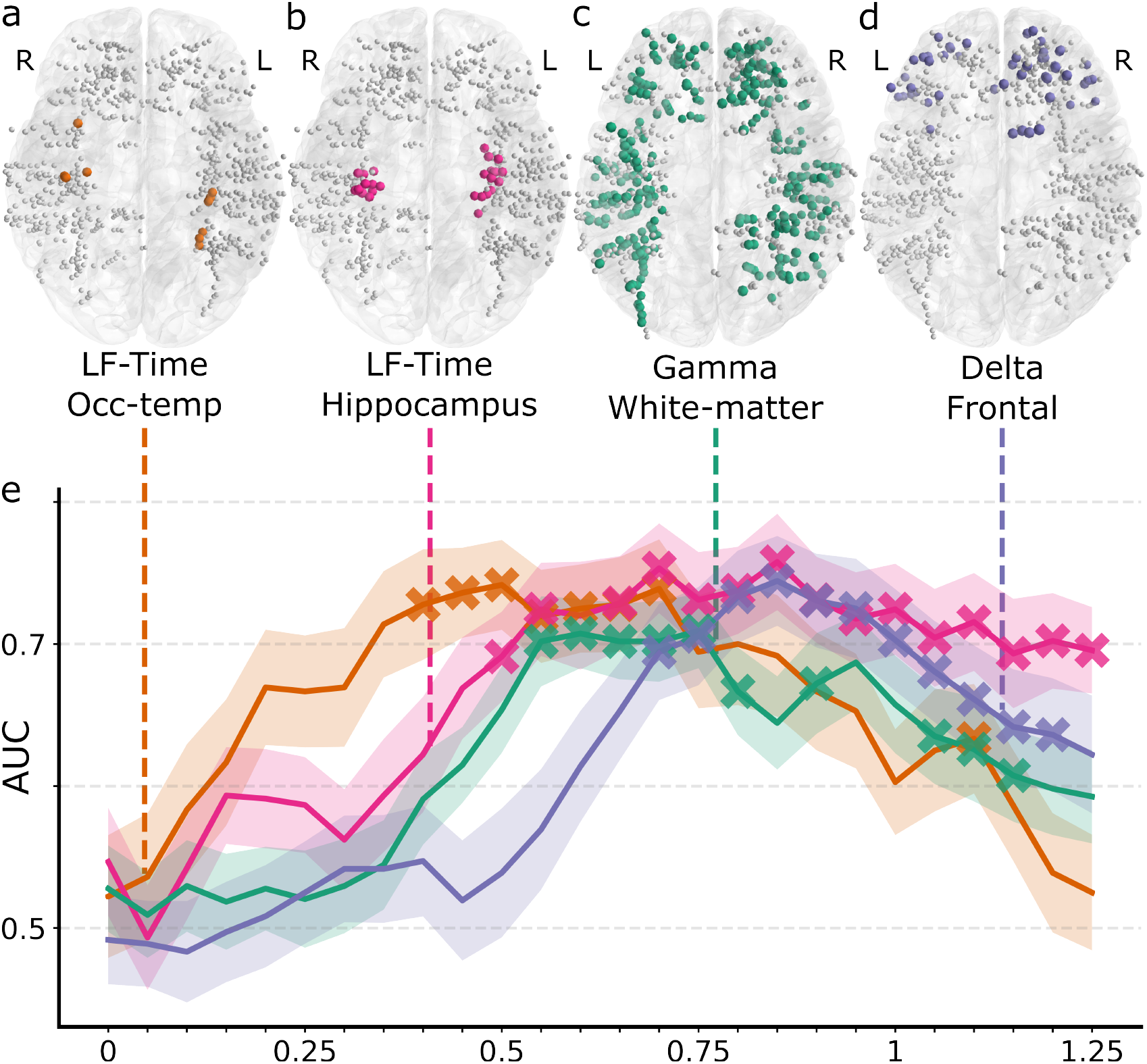
(e) Mean prediction AUC after stimulus presentation of selected regions of interests and features over time to highlight the temporal dynamics of different decision-making processes related to different levels of available information. Standard error for each time point is highlighted by the shaded area around each line plot. Time domain occipital-temporal sulcus (a), time domain Hippocampus (b), gamma white-matter (c) and delta frontal lobe (d) feature locations visualized in joint MNI space. Significance as determined by a mixed-effects model controlling for participant variance is highlighted with a x.

### Insular processes encoding the degree of available information prior to button press

Aligned to button press, gamma power features showed a particularly high decodability of informed vs. uniformed decision-making with a mean AUC of 72% (std=0.07; b=0.22, SE=0.03, z=6.97, p*<*0.0001, 95% CI=[0.16, 0.28]). To a lesser degree decoding from alpha power features also resulted in significant classification of information level with a mean AUC of 65% (std=0.06; b=0.15, SE=0.03, z=4.72, p*<*0.0001, 95% CI=[0.09, 0.21]). Delta fatures with a mean AUC of 58% (std=0.09; b=0.09, SE=0.04, z=2.11, p=0.02 (n.s. after bonferroni correction), 95% CI= [0.01, 0.16]), theta features with mean AUC of 55% (std=0.07; b=0.05, SE=0.03, z=1.55, p=n.s., 95% CI [−0.01, 0.11]), beta features with mean AUC of 57% (std=0.13; b=0.08, SE=0.06, z=1.26, p=n.s., 95% CI=[−0.04, 0.19]) and LF time domain features with a mean AUC of 57% (std=0.12; b=0.07, SE=0.05, z=1.38, p=n.s., 95% CI=[−0.03, 0.18]) did not achieve a decoding AUC significantly above 50% (also see Fig. 2). A further investigation of individual time intervals and regions of interest (ROIs) highlighted the importance of gamma power in insular regions in decoding information level (see Fig. 4). Based on this result, we performed an additional exploratory analysis to further characterize the insula subregion relevant for decoding informed and uninformed decisions by splitting contacts located in the insula into anterior and posterior insular contacts. Anterior and posterior insula contacts were determined using MNI labels of the respective contacts as well as their location related to the posterior outline of the short gyri of the insula. Seperate analysis of the decodability of information level from anterior and posterior insula gamma features showed higher decodability with more significant time points from anterior contacts. Furthermore, to localize the insular channels most predictive of information levels, we used single-channel decoding and one-sided t-tests to check whether the corresponding decoding AUC was higher than chance level. As depicted in Fig. 4 c, channels with the highest t-values were located in anterior insula and the distribution of channels with t-values above 2 was skewed towards the anterior part of the insula. Based on both analyses, we hypothesize that the location of relevant neural processes related to available information coding is predominantly in anterior insula regions.

**Figure 4.**
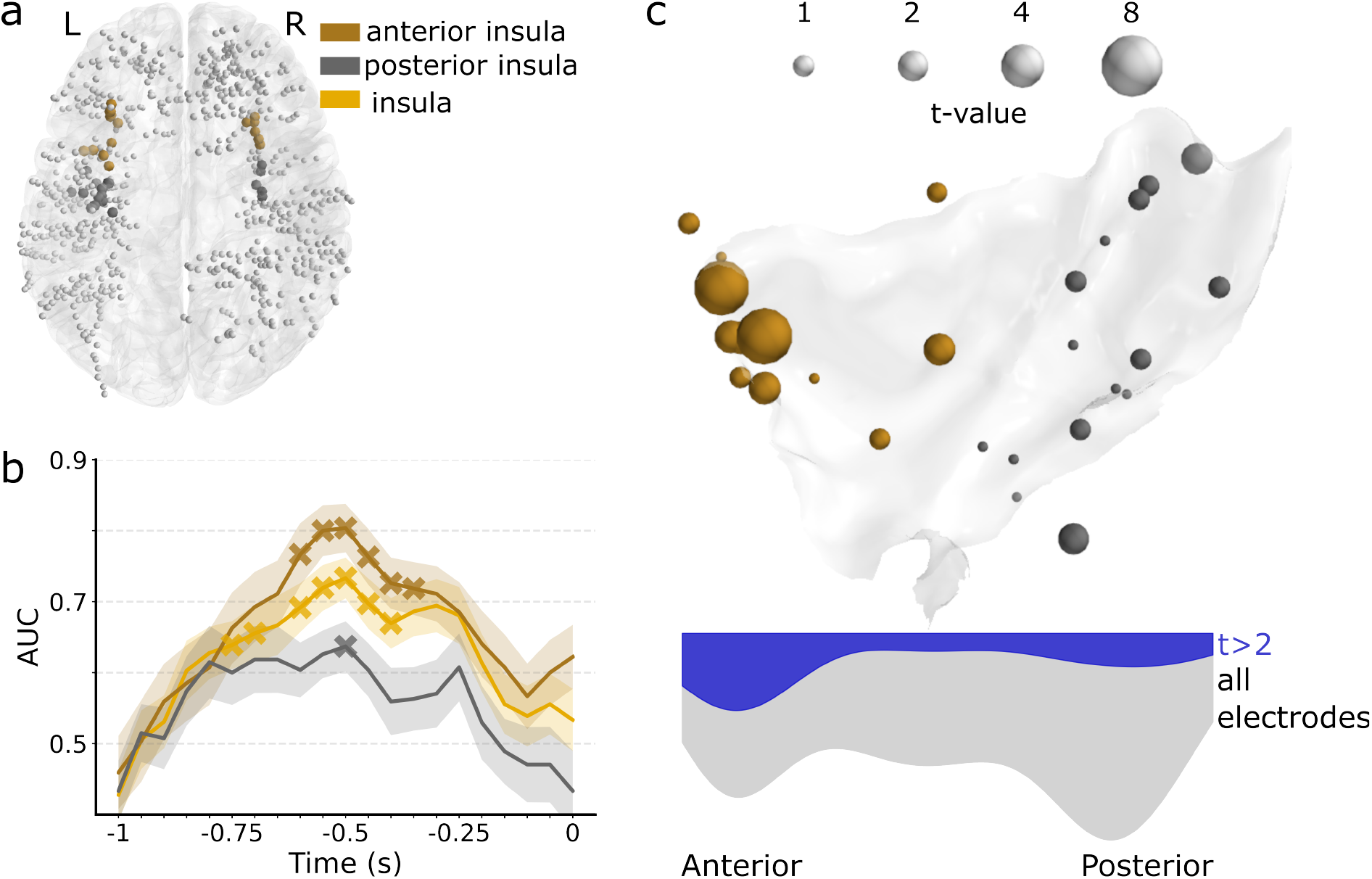
Insular involvement in coding available information: a) Locations of all electrodes located in insula and split into anterior and posterior insula. b) Decoding AUC for gamma features recorded from insular subregions over time. Standard error for each time point is highlighted by the shaded area. Significant decoding above chance level is indicated by a ‘x’. c) T-values of single channel decoding results plotted on the right insula MNI shape. Channels located in the left insula are mirrored. Distribution of all channels and channels with a t-value above 2 (blue) are plotted on a scale from anterior to posterior.

## Discussion

### High predictability of informed vs. uninformed decision-making

Our results highlight that decoding informed versus uninformed decision-making is possible from various time intervals both after stimulus onset and before button press. A maximum mean decoding AUC of 79% was achieved after stimulus onset and a mean decoding AUC of 72% was obtained in the time interval before button press. Band-pass filtered low frequency features aligned to stimulus onset yielded the highest AUC. This feature has also been previously identified as an important feature by Sadras et al. and Valeriani et al.^24,25^. Our results particularly replicate the EEG-study results by Sadras et al. showing a slightly higher mean decoding AUC of this feature using a similar decoding pipeline (79% compared to 75%)^25^. Similarly to their results, we do not find low frequency time domain features to be significantly predictive when aligned to participant response. Significant decodability of information level from gamma features is in line with previous results from Castelhano et al.^26^. They apply a task design comparable to the design in the study by Sadras et al. where decision-making based on scrambled vs. clearly visible objects (which can be interpreted as higher vs. lower available information level) is analyzed. They show related differences in gamma band power both aligned to stimulus onset as well as to participant response which we could replicate in our analysis^26^.

### From visual processing to cognitive control

The temporal dynamics of predictability of informed vs. uninformed decision-making from relevant regions of interest reveals further insights into relevant involved brain regions and underlying neural processes. Early decodability of information level approximately 400ms after stimulus onset was achieved from electrodes located in the temporal-occipital sulcus. This sulcus spanning from occipital lobe along the temporal lobe has been previously identified as a visual processing area for example connected to color processing^27^ and the connection of visual patterns to a phonological representation^28^. This could directly relate to the participants perceiving the different stimuli and potentially trying to name them as part of a memory strategy. Subsequently, we observe decodability of information level from stimulus-locked white matter gamma activity. Previous literature identified Gamma activity in visual cortex related to decision-making at different information levels to be linked to visual processing^26,29^. In this study we can not directly test decodability of informed and uninformed decisions from visual cortex contacts, as electrode sampling in our study is not sufficient for testing the significance of occipital regions across multiple subjects.

However, decodability of information level from gamma power in white matter contacts could indirectly relate to visual processing. White matter activity is hypothesized to be caused by communication between different brain regions or could originate from activity spreading from adjacent brain regions^30^. This could indicate a relatively early predictablility of the degree of available information related to neural activity during visual processing spreading from visual cortex. A slightly earlier timing than for white matter gamma is also observed for decodability from low-frequency time-domain features in Hippocampus. Hippocampus is known to play an important role in memory formation and memory retrieval particularly when making decisions informed by past experiences^31^. Memory retrieval could, according to our results, be a continuous process during decision-making lasting over a prolonged time window. Lastly, we also observe late decodability of available information level from frontal delta power features. Previous studies have linked delta oscillations in frontal lobe to the inhibition of interference for increasing task performance^32^ particularly during decision-making^33^. Delta oscillations can in this sense be understood as a modulator of networks related to either attention to external stimuli or internal concentration^32^. Therefore, differences in delta power between informed and uninformed decision-making could be hypothesized to be related to a varying necessity of focusing on the stimuli e.g. to perceive new stimuli versus internal concentration needed to remember information from previous trials. In summary, our results show that differences between informed and uninformed decision-making can be reflected in multiple neural processes starting with visual processing over memory retrieval to higher level control mechanisms, following a clear processing timeline.

### Anterior insula as a decision-making hub

Related to button-press, we find gamma band power features in insula electrodes to show a high decodability of informed versus uninformed decision-making. This is strongly in line with the findings by Goueytes et al. that identify insular channel gamma activity to reflect evidence accumalation over time^34^. Furthermore, Castelhano et al. localize decision-related lower gamma power, that differs between trials with higher or lower levels of available information, in decision network regions and particularly the anterior part of the insula^26^. Additional analysis separating insular electrodes into anterior and posterior insula indicate that decodability of degree of available information in the insula is mainly driven by anterior insula electrodes. The important role of anterior insula during decision-making has been described in previous literature^35–37^, linking it to a variety of decision-making processes including uncertainty during decision-making^38^. The posterior insula on the other hand has been described to be predominantly involved during the evaluation phase taking over a more general role in sensorimotor processing^35^.

### Coding of available information as a distinct neural construct

By predicting informed from uninformed trials, we evaluate the hypothesis that different levels of available information are coded differently in underlying neural processes. However, neural differences between first and third stimulus presentation could also be hypothesized to be caused by other differences between informed and uninformed trials. In this section, we want to further distinguish available information coding from other partially overlapping cognitive concepts and argue that the decoding of different information levels in our analysis indeed identifies neural processes that are distinct from novelty detection, reaction time decoding, and the neural representation of decision-confidence.

Differences in neural signatures related to the processing of novel vs. familiar stimuli have been commonly described in event related potential differences within the first 500ms after stimulus onset^39–41^. Our decoding results, however, capture not only low-level visual processing, which would be hypothesized to peak around 300ms after stimulus onset. This is demonstrated by significant decoding performance that consistently extends past the first 500ms after stimulus onset for all the relevant regions of interest. On top of that, we show that decoding is also possible when aligning neural features to button press, which leads to a distortion of stimulus-related neural activities due to varying reaction times.

Furthermore, while reaction times are significantly faster during informed decision-making on a group level, these timing differences can not explain our classification results. While decodability is consistent across participants, differences in reaction time vary between participants. Specifically, for participants P02 and P03, that show average response time differences close to zero, the classification AUC across features is comparable to the classification results achieved for other participants exhibiting bigger timing differences (see Fig. 2).

Lastly, we want to differentiate the cognitive concept of available information coding from decision confidence. In our task-design, participants do not report subjective decision-confidence but we use an objective prediction target of whether stimulus information was previously provided during the task. Available information and decision confidence inherently show an overlap, as they are both expected to correlate with improved decision-making performance (significant improvement of decision accuracy during informed decision-making is shown in Figure 1) and those correlations are also established in previous literature^24,34^. In previous literature, the term decision-confidence has sometimes been used to also describe the decoding of available information^25^. However, Goueytes et al.^34^ clearly establish a difference in neural processes between information accumulation and the coding of self-reported decision-confidence showing representations of the two processes both in overlapping as well as distinct brain regions. Furthermore, it is particularly the decoding of objective measures, such as the coding of available information, that is relevant for establishing a decision-making BCI to improve decision-making quality by prompting additional information seeking behavior. Decision-confidence, while correlated with performance, can be misplaced as it can, as previously established, be influenced by external factors^4^ as well as impacted by psychiatric, developmental and neurodegenerative disorders^8–11^. Valeriani and colleagues show that BCI predicted confidence is indeed a better predictor of participant accuracy than subjective decision confidence, highlighting the potential of enhancing decision performance based on neural markers^24^. While this is an exciting finding, we consider decoding the current information level as superior to predicting whether a user will give a correct response for usage in a BCI. We hypothesize that prediction of available information can generalize better to new situations with multiple or ambiguous response options, while still capturing the direct tradeoff between time needed for additional information seeking and decision-making quality^1^. This hypothesis should however be tested in future research by directly evaluating the potential of improving decision-making based on feedback predicted from different cognitive features in a variety of decision-making tasks and situations.

## Conclusion

Our results identify multiple high-performing temporal and oscillatory features for significantly differentiating between informed and uninformed decision-making. In particular, gamma, low-frequency time-domain and delta features can be related to relevant underlying neural processes. Indeed, our decoding results, both aligned to stimulus onset as well as to button press, in both cortical as well as regions of interest located below the cortical surface, highlight the complex dynamics of decision-making processes that encode the availability of decision-related information. The confirmation of prior results from previous studies, investigating the degree of available information using picture modifications, further validates the robustness of our decoding results and confirms that we are indeed capturing levels of information during decision-making as a general mechanism consistent across task designs^25^. However, we extend on previous results obtained from EEG analysis with the inclusion of deep brain regions and highlight the important role of regions such as hippocampus, occipital-temporal sulcus and insula sampled only by sEEG electrodes. Future research could evaluate further brain regions implied in decision-making for their relevance in coding available information not sufficiently sampled in our study. Of particular interest might be anterior cingulate cortex^42^, amygdala^43^ as well as the basal ganglia^44^. While our results provide insights into the cognitive processes and their temporal dynamics linked to neural information coding, we see additional relevance for our results to provide a foundation for future research investigating or implementing decision-making BCIs. Although our analyses only include offline results in a artificial decision-making task, they also provide fundamental information about different temporal dynamics of decision-making and regions of interest that can be helpful for translation into a closed-loop decoding setup. Through the selection and combination of previously identified time intervals and features, we expect the possibility to individualize the BCI to study participants to ultimately further enhance the prediction performance. We further think that it is of high importance to evaluate such a decision-making BCI using real-time decoding in a variety of tasks including situations of daily life as well as different user populations. Ultimately, we envision an application that can greatly improve decision-making performance and potentially improve quality of life for patient populations suffering from the effects of impaired decision-making.

## Methods

### Participants and Data recording

Six participants, between 21 and 52 years old (2 female, mean age=34.3 years, std=9.35), previously diagnosed with medication resistant epilepsy, were included in the study. All participants had previoulsy underwent implantation of platinum–iridium stereotactic-electroencephalography (sEEG) electrodes (Microdeep intracerebral electrodes; Dixi Medical, son, France) for invasive inpatient monitoring prior to resection surgery. Each electrode contained 5–18 contacts (2 mm long, 0.8 mm in diameter and 1.5 mm inter-contact distance). Two stacked Micromed SD LTM amplifiers (Micromed, S.p.A., Treviso, Italy) were used to record the neurophysiological local field potential (LFP) data. One contact located in white matter that was previously determined to not show any epileptic activity was used as a reference during recording. LFP data and behavioral markers were recorded simultaneously and saved using lab streaming layer^45^. Electrode implantation locations were determined only based on clinical considerations and all experiments were conducted under the supervision of healthcare professionals.

Participants volunteered to participate in the research and signed an informed consent form. The study was approved by the review board of Maastricht University and Epilepsy Center Kempenhaeghe (METC 20180451).

### Electrode localization

Electrodes were spread over wide parts of the brain, providing a coverage of both cortical and subcortical regions (Fig. 1 b). To determine the anatomical location of electrode contacts, we followed the same pipeline as described in previous work^46^, co-registering anatomical T1-weighted MRI scans to post-implantation CT scans. Using Freesurfer, the MRI was parcellated and anatomical locations were determined based on the Destrieux atlas. To depict the electrode locations of all subjects together, the brains and electrode locations were warped to MNI152 space.

### Task

Participants performed a decision-making task consisting of 90 trials divided into 10 blocks (compare^47^). During each trial, a fixation cross was shown for 500ms, followed by a one second presentation of a visual stimulus (marked as event “Stimulus Onset” in Fig. 1). The stimuli consisted of pictures randomly selected from the Novel Object and Unusual Name (NOUN) Database^48^. After stimulus presentation, participants were instructed to make a decision by pressing either the left or the right arrow button on a keyboard (also referred to as the event “Button Press”). This way they indicated to which of two groups (left or right) they thought the stimulus belonged to. After button press, a fixation cross was presented for 500ms. Subsequently, feedback was given using color and sound that indicated a correct or incorrect decision. During each block, three different stimuli were presented 3 times each. The order in which stimuli were presented was randomized. Stimuli were randomly, but consistently for all three presentations, assigned to one of two groups (left or right). That meant that while in the first presentation trial the participants had to guess which group a stimulus belonged to, during the second and third presentation trial, they could base their decision on previously obtained information (also see Fig. 1 a) for an overview of the task design). Before the start of the task, participants first conducted a training block to familiarize themselves with the task design.

### Data processing

Recorded electrophysiological data was re-referenced to the common average of each electrode shaft. Both, different oscillatory and low frequency domain features were considered (Fig. 2 a) in the further analysis. Delta (1-3Hz), theta (4-7Hz), alpha (8-12Hz), beta (13-30Hz), gamma (30-100Hz) frequency band activity was extracted by filtering the raw neural time series using a bandpass filter (butterworth, order=3). To compute the band power of each frequency band, the Hilbert transform was applied and the envelope of the Hilbert transform was computed by taking the absolute of the transformed signal.

The time domain information in the low frequency range was retrieved by applying a band pass filter (butterworth, order=3) between 1 and 40Hz.

In a consecutive step, we split the data into trials aligned to one of the two task events (Stimulus Onset or Button Press). Trials were split into multiple time windows with a length of 500ms and were computed using a sliding window. The first time window started at the onset of the respective task events capturing 0 to 500ms after Stimulus onset or accordingly −500 to 0ms before button press. Afterwards, the sliding window was shifted with a frameshift of 50ms forwards in time until 1 second after Stimulus onset and backwards in time until 1 second before button press. Consequently, there were 11 partially overlapping time windows extracted for both events capturing one second of recording time each (also compare Fig. 2). To extract the oscillatory features of each trial, the average of the respective frequency envelope (delta, theta, alpha, beta, gamma) within each time window was computed for every channel. Time-domain features of each trial were extracted by downsampling the low-pass filtered time domain signal to 8Hz, thus resulting in 4 datapoints per trial, time window and channel.

### Decoding pipeline

Labels of information level were assigned to the respective trials based on task information. In trials in which a visual stimulus was presented for the first time, we assumed that the participant was not able to leverage previous information and was therefore forced to make an uninformed decision. On the other hand, in a trial in which a visual stimulus was presented for the third time the participant could exploit information from previous trials and make an informed decision. The trials were labeled as uninformed versus informed decision-making trials accordingly.

In our machine-learning pipeline, we used a min-max scaler and a logistic-regression classifier to test the predictability of the previously extracted oscillatory and temporal features in a 10-fold cross validation for differentiating between the two classes. To evaluate the performance of the classification, we utilized the receiver operating characteristics area under the curve (roc-AUC) score (also referred to as AUC). To test whether the AUC significantly exceeds chance level, we used a one-sample, one-sided linear mixed-effects model on the results. In this approach, the AUC scores of each fold of the cross-validation were considered as the fixed effect predictor and participant number was treated as a random effect. With this approach, we tested for a performance significantly above 50% while controlling for variability between subjects, thus ensuring that the effect is consistent across subjects. Correction for multiple testing was performed using Bonferroni correction.

During the analysis, we used the machine-learning pipeline in two consecutive analysis steps. In a first step, we evaluated the overall decoding performance per frequency band. The intention was to show the general feasibility of decoding informed versus uninformed decision-making aligned to the two events from the selected features. Therefore, we combined the information in all channels for all extracted time windows per event, one second after stimulus onset and one second before button press accordingly. We used this high-dimensional feature vector to classify informed and uninformed decision-making using each of the frequency bands separately.

In a second step, we zoomed in on specific time-windows and specific regions of interest to test their relevance for distinguishing between informed and uninformed decision-making. We selected the occipital-temporal sulcus, hippocampus, frontal lobe and insula as regions of interest, as they have previously been associated with decision-making processes and were sampled in at least 4 individual participants. White-matter contacts were also included as regions in the analysis to test for neural activity not originating from the included regions of interest. Therefore, we focused on the temporal dynamics per frequency band in 500ms time windows individually using the previously described prediction pipeline. Additional time windows nefore and after stimulus onset as well as button press were added to better visualize the temporal dynamics. The insula region was further split into smaller subregions using MNI labels as well as the outline of the short gyri of the insula to seperate contacts into anterior and posterior insula for decoding informed versus uninformed decision-making. Single channel decoding using the same classifier was performed for insular channels using all time-windows located in the relevant time window. Significance was tested using a one-sided t-test and t-values are extracted.

## Author contributions statement

L.M., S.A.H, M.L.F.J and C.H. designed the task. M.V., J.P.D. and S.T. recorded the patient data. S.G. performed the analysis and wrote the first draft of the manuscript. P.W., M.L.F.J, P.L.K and Y.T. provided valuable input and feedback during the project.

## Acknowledgements

P.W., P.L.K., M.V. and C.H. acknowledge funding by the project INTENSE (with project number 17619 of the research programme NWO Crossover Programme) which is (partly) financed by the Dutch Research Council (NWO). C.H. acknowledges funding by the Kavli Foundation.

## Data and Code availability

De-identified data is publicly available at https://doi.org/10.17605/OSF.IO/PEX8R. The corresponding Python code is shared on GitHub and is available at https://github.com/GimpleSophia/Decoding-brain-wide-signatures-of-uninformed-choices-for-BCI-assisted-decision-making.

